# Hinokitiol alters Gene Expression in *Aspergillus fumigatus*, protects against fungal keratitis by Reducing Fungal Load, LOX-1, Proinflammatory cytokines and Neutrophil Infiltration

**DOI:** 10.1101/2022.02.18.480984

**Authors:** Shuqin Ma, Hao Lin, Xudong Peng, Cui Li, Qian Wang, Qiang Xu, Mengting He, Dan Shao, Xing Liu, Guiqiu Zhao, Jing Lin

**Author notes:** **Co-correspondence authors:** Jing Lin and Guiqiu Zhao Jing Lin,; Guiqiu Zhao. Co-first authors: Shuqin Ma and Hao Lin.

## Abstract

Hinokitiol (HK), a tropolone-related compound found in cupressaceous plants, has antifungal and anti-inflammatory properties. But its roles in *Aspergillus fumigatus* ( *A.F.* ) keratitis remains unknown. This study examined the antifungal activity of HK against *A.F.* and investigated its possible mechanisms at the ultrastructural and transcriptional levels. We tested the minimum inhibitory concentration (MIC) of HK against *A.F.*. Our data showed that the MIC50 was 2 μg/ml and MIC90 was 8 μg/ml, and HK could significantly inhibit the spore germination and mycelial growth of *A.F.*. Scanning electron microscopy (SEM) and transmission electron microscopy (TEM) observations showed that HK induced significant changes in hyphal morphology and microstructure, including cell membrane rupture and intracellular structural disorder. Adhesion assay and biofilm assay experiments showed that HK reduced the adhesion of spores to human corneal epithelial cells (HCECs) and inhibited the formation of biofilms. The genetic changes of *A.F.* after HK treatment were analyzed by RNA sequencing, and a total of 2487 differentially expressed genes (DEGs) were identified, including the down-regulated DEGs related to carbohydrate and various amino acid metabolism, ribosome biogenesis, asexual reproduction, and up-regulated DEGs related to steroid biosynthesis through GO and KEGG enrichment analysis. RNA-seq results suggested that HK may play an antifungal effect by destroying the structural integrity of its wall and membrane, inhibiting protein biosynthesis and the growth and reproduction of *A.F*..10 μg/ml HK, which had no effect on proliferation and migration of HCECs, was used in the treatment of murine fungal keratitis (FK). We found that HK eyedrop treatment could reduce the severity of FK by reducing corneal fungal burden, neutrophil infiltration, and the expression of LOX-1 and pro-inflammatory factors. HK is expected to be a safe and effective new drug for FK treatment.

**Author summary:** FK is a serious common blind-causing eye disease. The current clinical antifungal drugs have the disadvantages of low bioavailability and high corneal toxicity, which greatly limit the clinical effectiveness. Therefore, there is an urgent need for a new, safe and effective treatment for FK. HK had shown great antifungal capacity against *A.F.*, amazingly well anti-inflammatory effect and real improvement in murine FK in our research. These findings provide a theoretical basis for the application of HK as an antifungal drug and reveal the potential of HK to be applied in the clinical treatment of FK.

## Introduction

FK is a serious infectious eye disease that can cause permanent vision loss. *A.F.* and *Fusarium* are common pathogenic fungi for FK in developing countries(1). The increase in plant-related ocular trauma, rational use of contact lenses, and irregular use of glucocorticoids all contributed to the increasing incidence (2–4). However, currently commonly used antifungal drugs such as natamycin and voriconazole eyedrops have shortcomings such as poor local penetration of the cornea, strong irritation, and high toxicity, and have obvious limitations in clinical application(4, 5). Therefore, it is of great significance to develop new safe and non-toxic antifungal treatment.

HK, also known as β-thujaplicin, is a natural tropone-related compound purified from cupressaceous plants(6). HK has a wide range of biochemical and pharmacological activities, and has been widely used in toothpaste and oral care gels with low cytotoxicity (7–10). Previous studies have shown that HK has a well antifungal effect, and inhibit the hyphal growth of *Candida albicans* (*C. albicans*) by blocking RAS signaling(11, 12). HK inhibits *Candida* biofilm formation at concentrations of 3.1-12.5 μg/ml and decreases mature biofilms between 12.5-400 μg/ml (12). Studies have also shown that HK has significant anti-inflammatory effects. In HCECs stimulated by polyinosinic acid, HK can inhibit the nucleus translocation of NF-κB and downregulate the expression of pro-inflammatory factors, protecting the cells from damages(13). In middle cerebral artery occlusion-induced thromboembolic stroke rats, HK can provide neuroprotection by inhibiting inflammatory responses and apoptosis (14).

The transcriptome is the collection of all transcripts produced by a species or a specific cell type(15). Transcriptome studies can study gene function and structure at the overall level, revealing specific biological processes and molecular mechanisms (16). Pan et al. revealed the molecular mechanism by which perillaldehyde induces cell death in *Aspergillus flavus* by inhibiting energy metabolism through transcriptome sequencing analysis (17). Tripathi et al. found that Puupehenone enhances the antifungal ability of antifungal drugs by disrupting Hsp90 activity and cell wall integrity pathways by transcriptome analysis(18). However, the effect of HK on the transcriptional profile of *A.F.* and the molecular mechanism remains unclear.

*A.F.* induces an innate immune response in FK (5). Pattern recognition receptors (PRRs) specifically recognize pathogen-associated molecular patterns (PAMPs) on fungal cell walls, activating signaling cascades consisting of neutrophils, macrophages, inflammatory factors, and chemokines(19). However, an excessive inflammatory response leads to the accumulation of large numbers of immune cells and cytotoxic substances, exacerbating tissue damages or delaying wound healing (20, 21). Inhibiting fungal burden and controlling excessive inflammation are considered as effective strategies to improve clinical outcomes of fungal infectious diseases. LOX-1, a lectin-like 52-kD type II low-density lipoprotein oxidizing membrane receptor, plays a pro-inflammatory role in FK in our recent studies(22, 23).

However, whether HK can exert anti-inflammatory and antifungal effects in FK and its possible mechanisms are still unclear. In this experiment, the effect of HK on *A.F.* was preliminarily verified in vitro, and its possible antifungal mechanism was studied at the microstructure and transcriptome levels. We also demonstrated the antifungal and anti-inflammatory effects of HK on the murine FK model and explored the underlying mechanisms. This study provides a new idea for clinical treatment of FK.

## Results

### HK inhibited the growth of *A.F*

To determine the antifungal effect of HK, we performed MIC experiments on *A.F..* The MIC data showed that HK inhibited the germination of *A.F.* spores and hyphae from 2 μg/ml, and inhibited 90% of the spore germination at 8 μg/ml (Fig. 1A, B). Time-kill test was used to evaluate the stability of HK against *A.F.*. Compared with the solvent control group, the absorbance value of the 2 μg/ml HK treatment group decreased significantly, and gradually decreased with the increase of concentration (Fig. 1C). When HK concentration was 8 μg/ml, the absorbance was close to 0, and the value did not increase with time extension within 96 hours, indicating that HK concentration of 8 μg/ml had a stable fungicidal effect on *A.F.*. In addition, *A.F.* was incubated with 0, 2, 4, 8 and 16 μg/ml HK or 0.1% DMSO and stained with Calcofluor white stain. Calcofluor white stain can bind to chitin in the fungal cell wall, giving it a bright blue color under fluorescent excitation. The results showed that with the increase of HK concentration, the number of *A.F.* hyphae decreased, the branches decreased, and the length shortened (Fig. 1D-I). With incubated with 16 µg/ml HK, *A.F.* hyphae and spores were not observed in the field (Fig. 1I).

**Fig 1.**
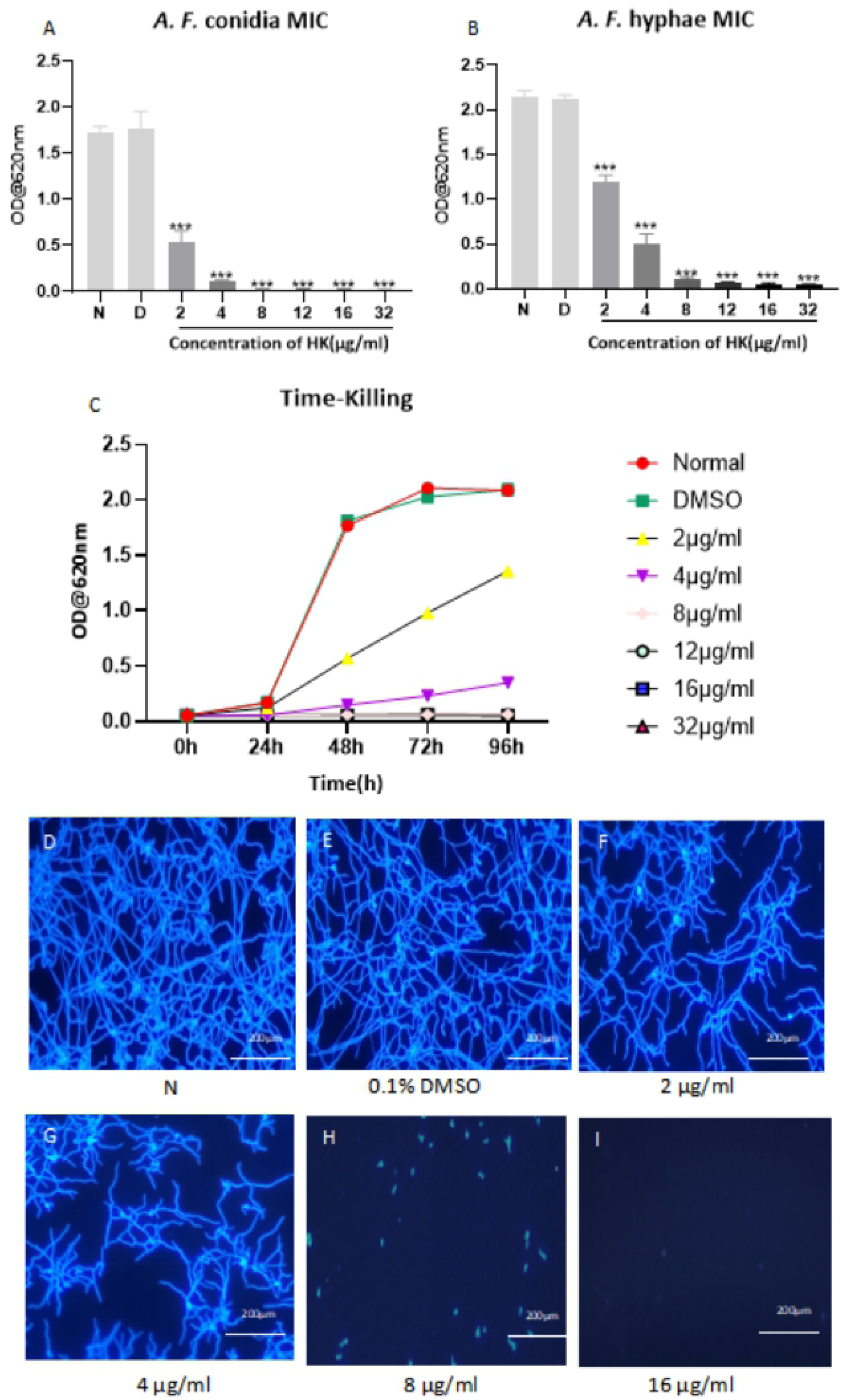
Antifungal effect of **HK** in vitro. *A.F.* conidia (A) or mycelium (B) were incubated with 2, 4, 8, 12, 16 and 32 µg/mL HK or 0.1% DMSO. Time-killing curves for *A.F.* exposed to 2, 4, 8, 12, 16, and 32 µg/mL of HK or 0.1% DMSO were performed over a period of 96 hours (C). HK decreased the fungal mass in a concentration-dependent manner at 36 has measured by Calcofluor white staining.

### HK inhibited *A.F.* through multiple mechanisms

We further explored the mechanism that how HK inhibited fungal growth using electron microscopy observations. The SEM results showed that the surface of *A.F.* in 0.1% DMSO medium was regular, uniform, smooth, without cracks, and in a normal growth state (Fig. 2A, B), while HK-treated hyphae had rough, distorted surfaces, forming pits and holes (Fig. 2C, D). PI staining was performed on hyphae to detect the effect of HK on the cell membrane permeability, which is not taken up by normal cells, but the cells whose membrane is disrupted. The fluorescence intensity increased with the increase of HK concentration, indicating that the effect of HK on fungal membrane permeability was dose-dependent (Fig. 2E, F, G). We used TEM to reveal the ultrastructural changes of the organism in response to HK. The control group was a typical ultrastructure of *A.F.*, with complete cell wall, membrane, distributed cytoplasm, regular nucleus, and the mitochondria were clearly visible, when other organelles were arranged in an order (Fig. 2H, I). However, the hyphal structures exposed to HK were highly deformed, with indistinct cell walls, discontinuous cell membranes, and disorganized internal structures. Their damaged cell membrane was completely separated from the cell wall, some organelles were missing, with few mitochondria remained (Fig. 2J, K).

**Fig 2.**
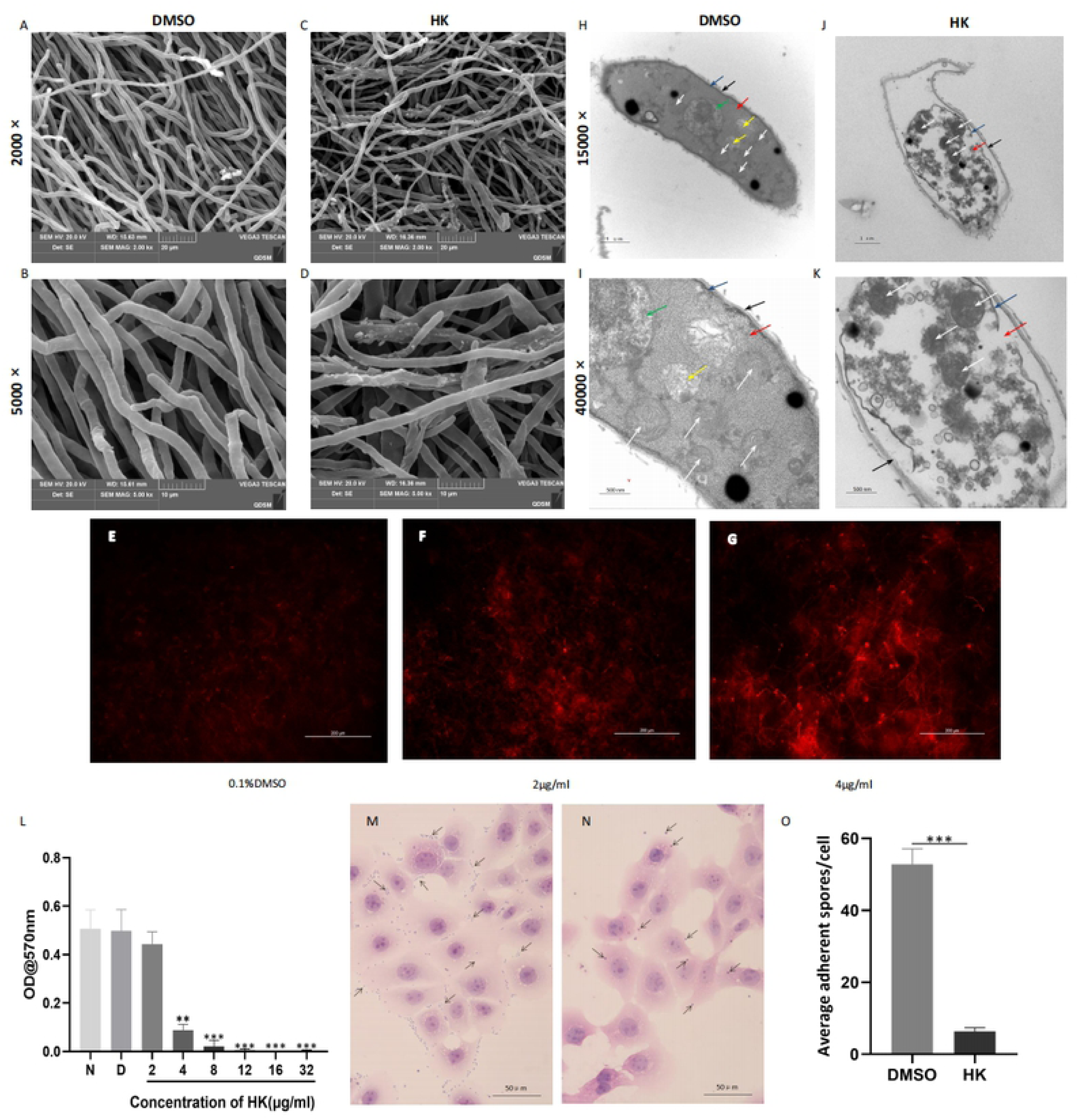
Multiple mechanisms of **HK** inhibited *A.F..* SEM images of *A.F* mycelium which were treated with 0.1%DMSO (A, bar=20µm; B, bar =IOµm) or 2µg/ml HK (C, bar= 20µm; D, bar= IOµm). PI staining showed compared with 0.1%DMSO group (E), cell membrane permeability increased after treatment with HK at 2 µg/ml (F) or 4 µg/ml (G; bar:I00 µm). The effect of HK at 4 µg/ml on the ultrastructure of *A.F.* was demonstrated by TEM (J, bar=lµm, K, <u>bar=500nm).</u> 0.1%DMSO group was used as normal control group (H, <u>bar=lµm,</u> I, <u>bar=500nm).</u> In TEM images, black arrows mark the cell wall, blue arrows mark the membrane, red arrows mark the cytoplasm, green arrows mark the nucleus, yellow arrows mark the endoplasmic reticulum, and white arrows mark the mitochondria. According to the absorbance values of crystal violet released from biofilm, HK significantly inhibited the formation of biofilm at 4 µg/ml (L). HCECs which were infected with *A.F.* condia were treated by HK (8 µg/ml, N, <u>bar=50µm)</u> or 0.1¾DMSO (M, <u>bar=50µm)</u> for 3 hours respectively. In the HE staining picture, black arrows indicated condia adhering to HCECs. Quantitative analysis was shown in figure O.

We also examined the effection about different concentrations of HK on fungal biofilm formation and the adhesion ability of spores. 4 μg/ml HK had a significant inhibitory effect on biofilm (Fig. 2L). The absorbance value decreased in a concentration-dependent manner. We tested the adhesion ability by counting the number of spores adhered to each HCEC. The number in HK group (8 μg/ml, Fig. 2N) was significantly reduced compared with the DMSO group (Fig. 2M). (Fig. 2O)

### RNA-Seq analysis

In order to explore the molecular mechanism of antifungal effect, samples from HK treatment group (Test group) and control group (Con group) with 36h incubation were tested by RNA-seq analysis. Table 1 shows all raw data and output statistics. The CleanBases of each library were above 6.1GB. In raw readings, the percent Q30 was over 94% in each group and the mapping percentage for each library was over 97%. The sequencing data was suitable for further biological analysis. 2487 DEGs were identified in the Test group, with 943 up-regulated genes and 1,544 down-regulated genes (Fig. 3A). Fig 3B showed the volcano plot analysis of all DEGs, and the DEGs we obtained have great difference and significance. To obtain a global view of gene expression profiles after exposure to HK, we clustered the differential genes to a heatmap (Fig. 3B), which represented the transcript levels of all DEGs between the two groups. The results showed that in the Test group, most of the DEGs exhibited significantly reduced expression patterns (blue bands), while in the Con group, most genes exhibited an up-regulated pattern (red bands). The three samples belonging to the Con group (Con1, Con2, and Con3) have similar expression patterns, as well as the three samples belonging to the Test group (Test1, Test2, and Test3), which proves that both the Con and Test groups have great sample repeatability.

**Fig 3.**
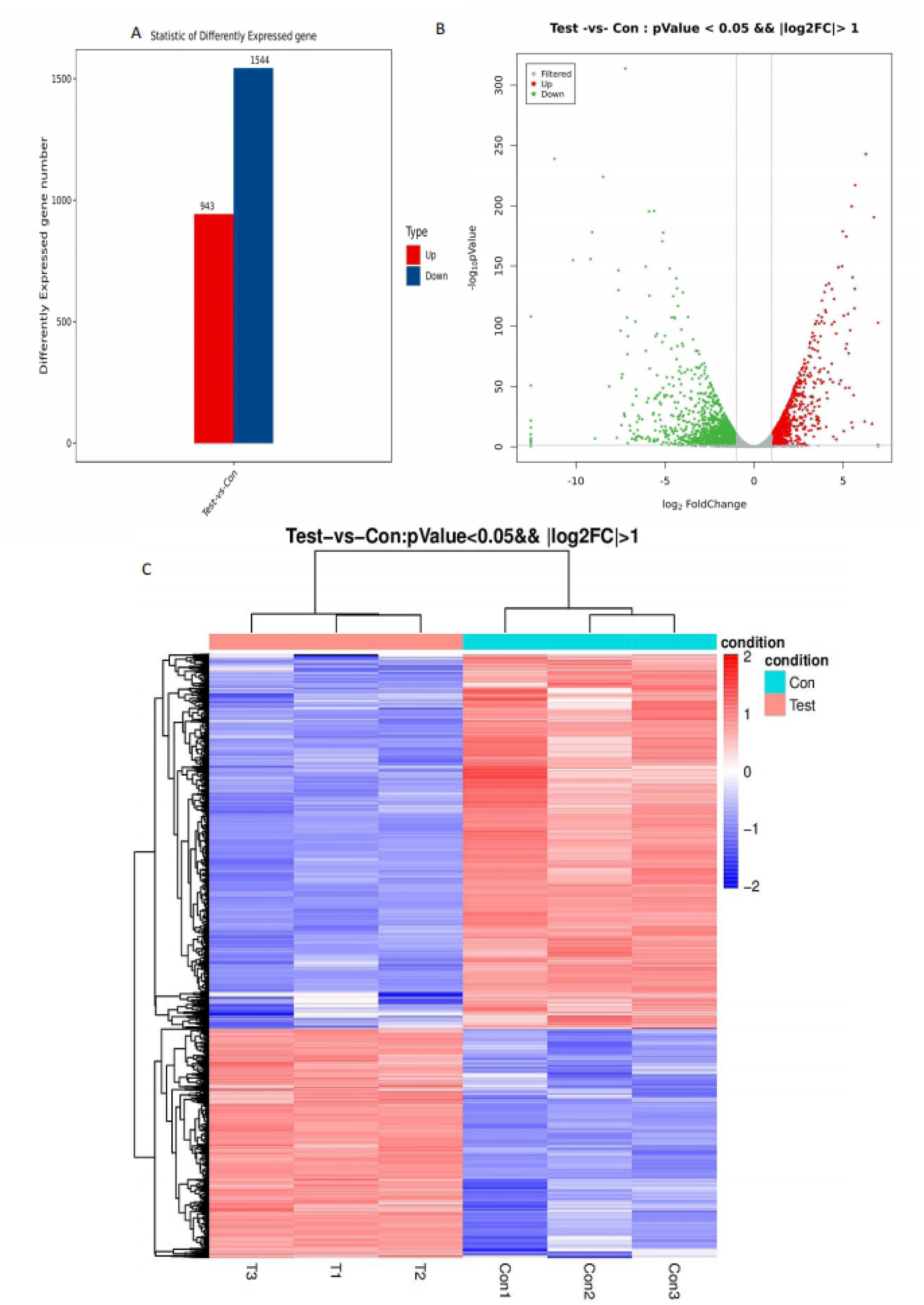
Transcriptional comparison in *A.F.* treated with 0.1%DMSO or HK (2µg/ml). Statistical histogram ofDEGs (A). Red bar represents up-regulated DEGs and blue bar represents down-regulated DEGs between HK treatment and the control. Volcano blot of DEGs (B). Up-regulated DEGs are shown in red dots and green dots represent down-regulated DEGs, while gray dots mark the genes with no significant difference. In the heat map, red represents the protein encoding gene with relatively high expression and blue represents the protein encoding gene with relatively low expression (C).

**Table I.** All raw data and output statistics of sequencing data.

| Sample | RawReads | RawBases | CleanReads | CleanBases | ValidBases | Q30 | GC | Mapped/% |
| --- | --- | --- | --- | --- | --- | --- | --- | --- |
| Con1 | 48.34M | 7.25G | 47.54M | 6.99G | 96.38% | 94.63% | 53.15% | 97.53 |
| Con2 | 49.96M | 7.49G | 49.47M | 7.33G | 97.83% | 95.94% | 53.49% | 98.23 |
| Con3 | 42.23M | 6.33G | 41.78M | 6.19G | 97.70% | 95.80% | 53.44% | 98.09 |
| T1 | 49.44M | 7.42G | 48.87M | 7.25G | 97.77% | 95.80% | 53.72% | 98.35 |
| T2 | 43.18M | 6.48G | 42.68M | 6.33G | 97.74% | 95.61% | 53.84% | 98.33 |
| T3 | 47.30M | 7.10G | 46.48M | 6.87G | 96.89% | 95.05% | 53.81% | 98.10 |

GO enrichment analysis was performed on DEGs with adjusted P value < 0.05, and their functions were described in combination with GO annotation results. All identified DEGs were classified into 64 subgroups (Fig. 4A). Among them, 23 subgroups belonged to biological process, 20 subgroups belonged to cellular component, and 21 subgroups belonged to molecular function. Cellular process, metabolic process and single-organism process were significantly distinct subcategories between Con and Test samples. In addition, there were certain DEGs mapped to localization, establishment of localization, cellular component organization or biogenesis, biological regulation and response to stimulus. In the cellular component, cell, cell part, organelle, organelle part, membrane, membrane part were the most significant enriched clusters. DEGs related to molecular function were mainly involved in the catalytic activity, binding and transporter activity subgroups (Fig. 4A).

**Fig 4.**
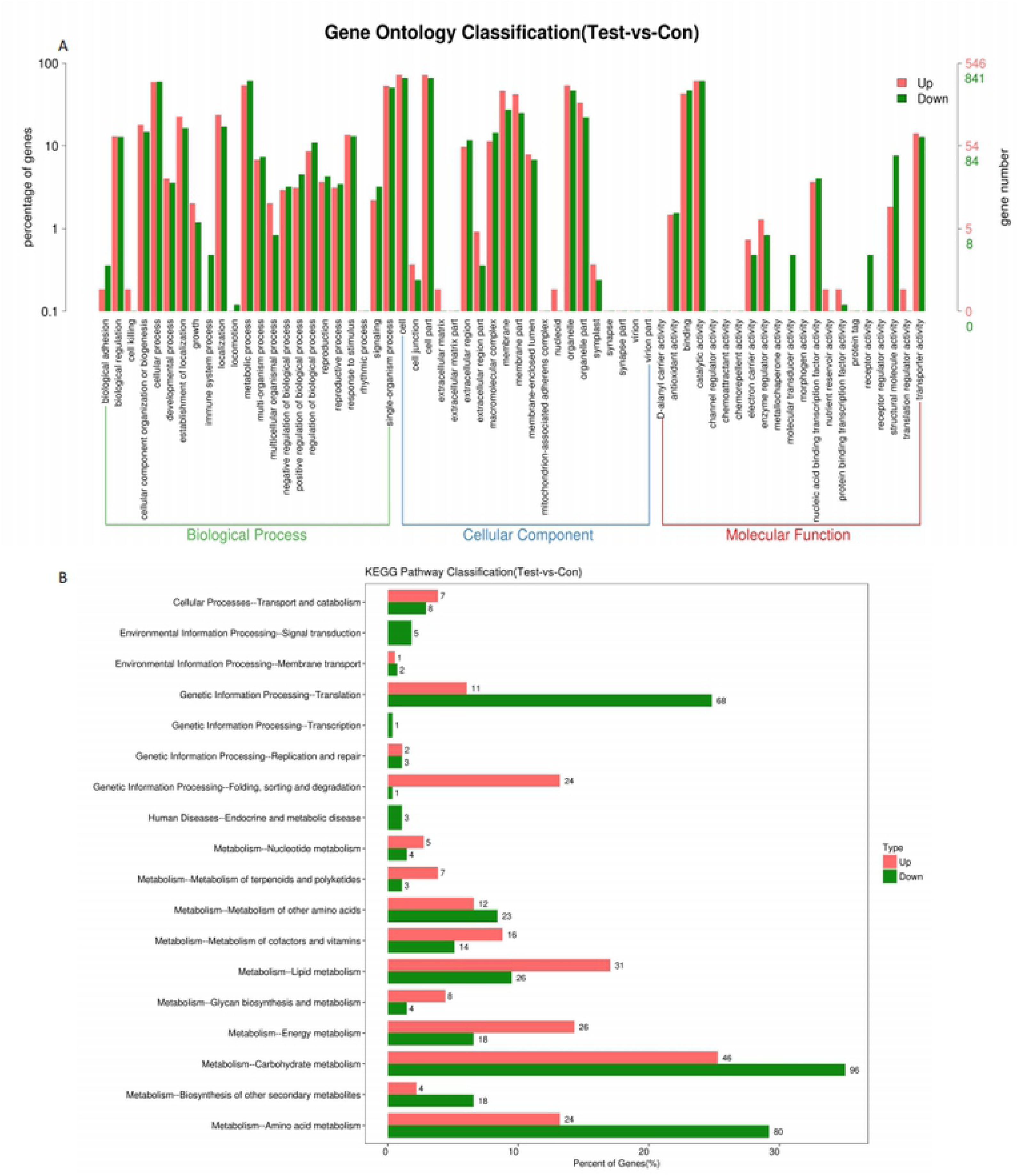
GO categorization and KEGG pathway categorization analysis of DEGs in *A.F.* treated with O.t¾DMSO or **HK** (2 µg/ml). Comparison of the distribution of up-regulated (red) and down-regulated (green) DEGs at GO categorization (A). KEGG pathway categorization distribution map of up-regulated (red) and down-regulated (green) DEGs (B).

Based on the KEGG pathway database, we aligned the identified DEGs with specific biochemical pathways. The results showed that these genes were mainly identified in carbohydrate metabolism, amino acid metabolism, genetic information processing-translation, lipid metabolism, genetic folding, sorting and degradation (Fig. 4B). KEGG enrichment analysis TOP20 pathways were mostly identified as ribosome and steroid biosynthesis (Fig. 5A). Among them, ribosome mainly showed a decrease, and steroid biosynthesis mainly showed an increase (Fig. 5B, C).

**Fig 5.**
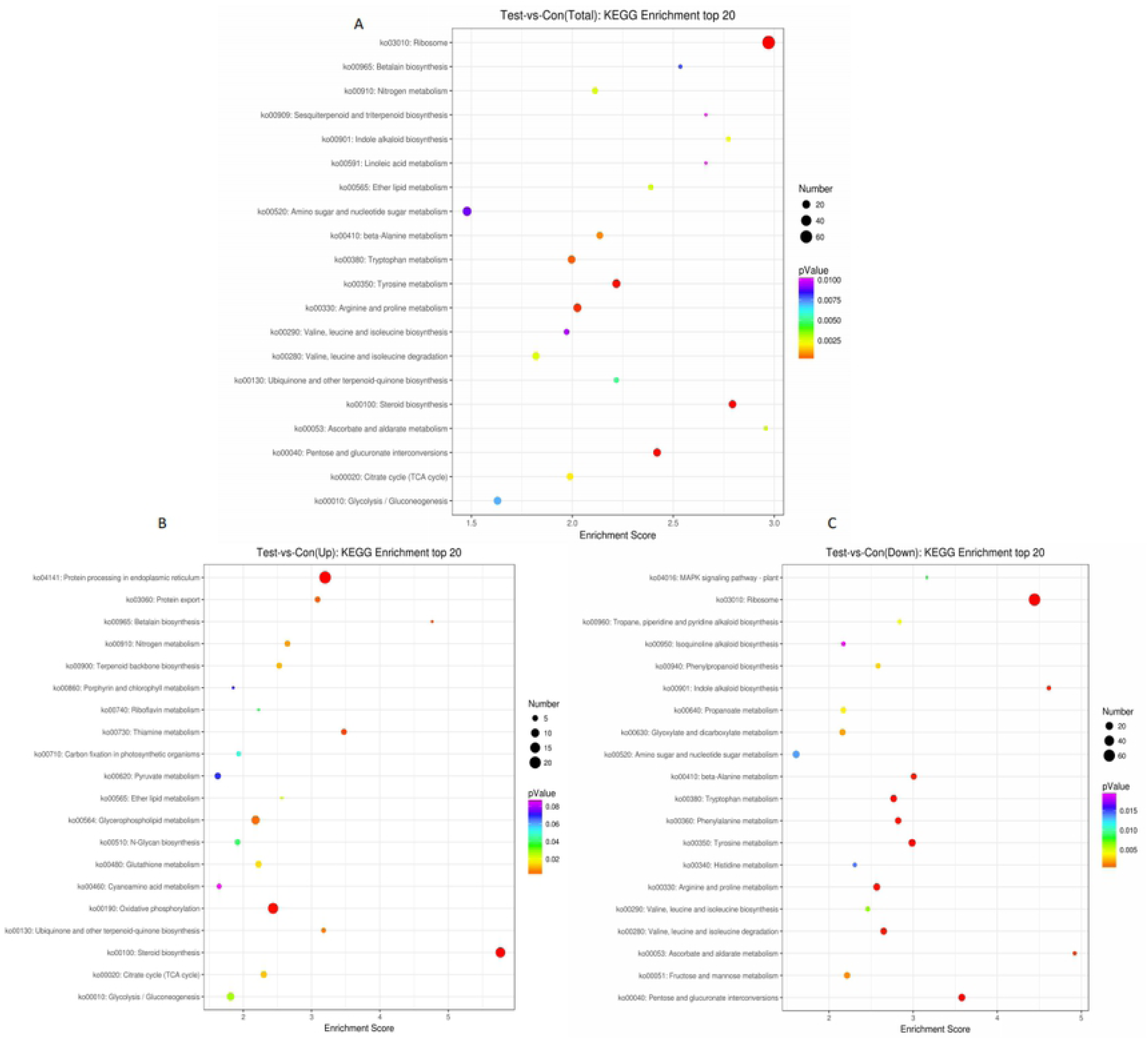
The top20 KEGG pathway categorization analysis of DEGs. Totle top 20 (A); Up top 20 (B); Down top 20 (C). The horizontal axis in the figure represents Enrichment. Items with larger bubbles contain more differentially encoded genes, and the bubble color varies from purple-blue-green to red, indicating the smaller enrichment pValue and greater significance.

### Validation of RNA-Seq

To verify the expression of RNA-seq genes, we picked 9 genes with different functions based on KEGG pathway analysis for qRT-PCR analysis. As shown in Figs 6A-6I, 8 selected genes were down-regulated and 1 was up-regulated. Although degree of change between each gene detected by qRT-PCR and RNA-Seq was different, the expression patterns of selected DEGs were very similar and showed a high correlation (R^2^=0.897) (Fig. 6J). These data suggested that RNA-seq analysis was reliable and accurate, and DEGs that may be involved in HK’s antifungal activity were shown in Table 2.

**Fig 6.**
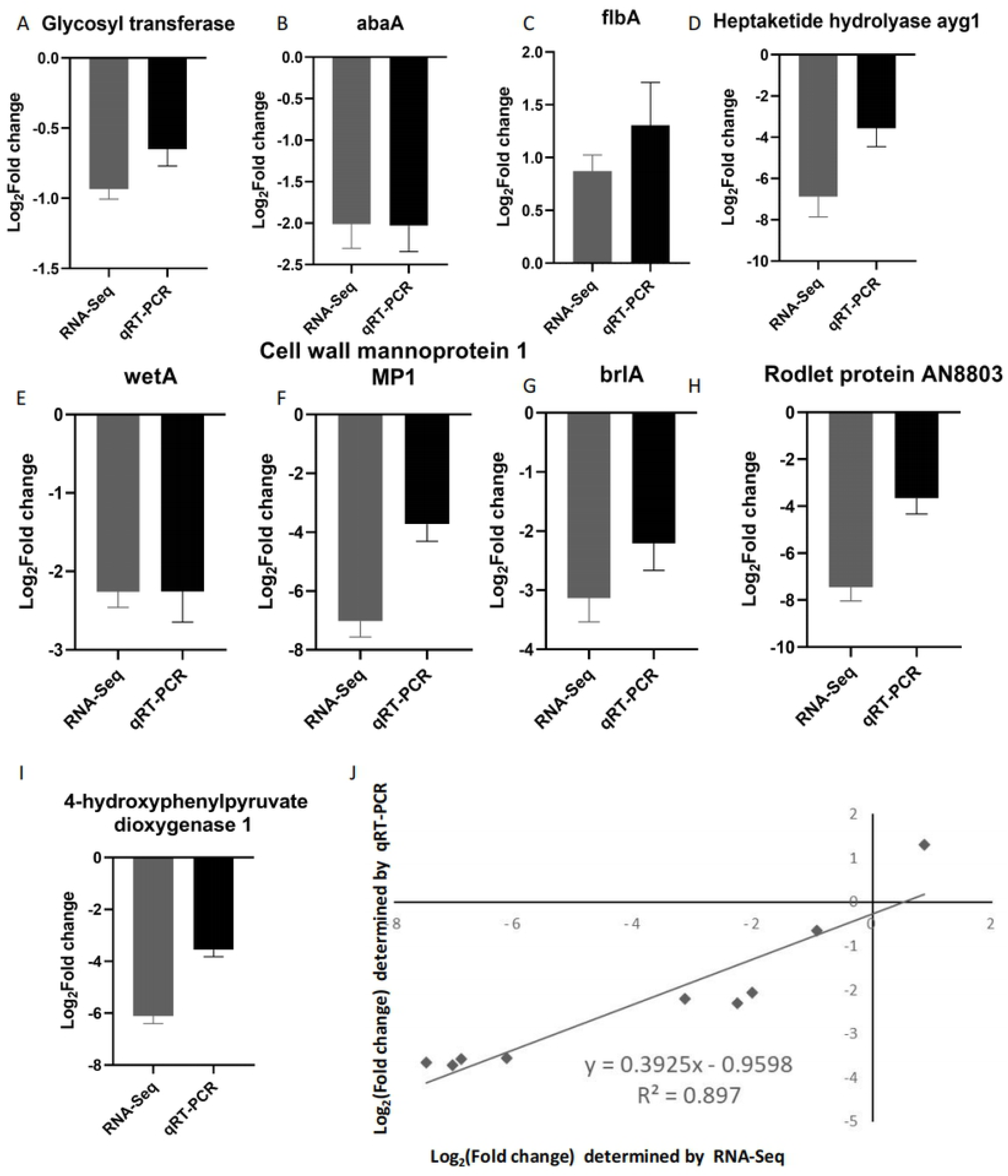
Correlation of expression level between RNA-Seq (gray) and qRT-PCR (black). The 9 selected genes were Glycosyl transferase (A), abaA (B), flbA (C), Heptaketide hydrolyase aygl (D), wetA (E), Cell wall mannoprotein I MP! (F), brlA (G), Rodlet protein AN8803 (H), 4-hydroxyphenylpyruvate dioxygenase I (I). Correlation between the RNA-Seq and qRT-PCR data are plotted in figure J.

**Table 2.**
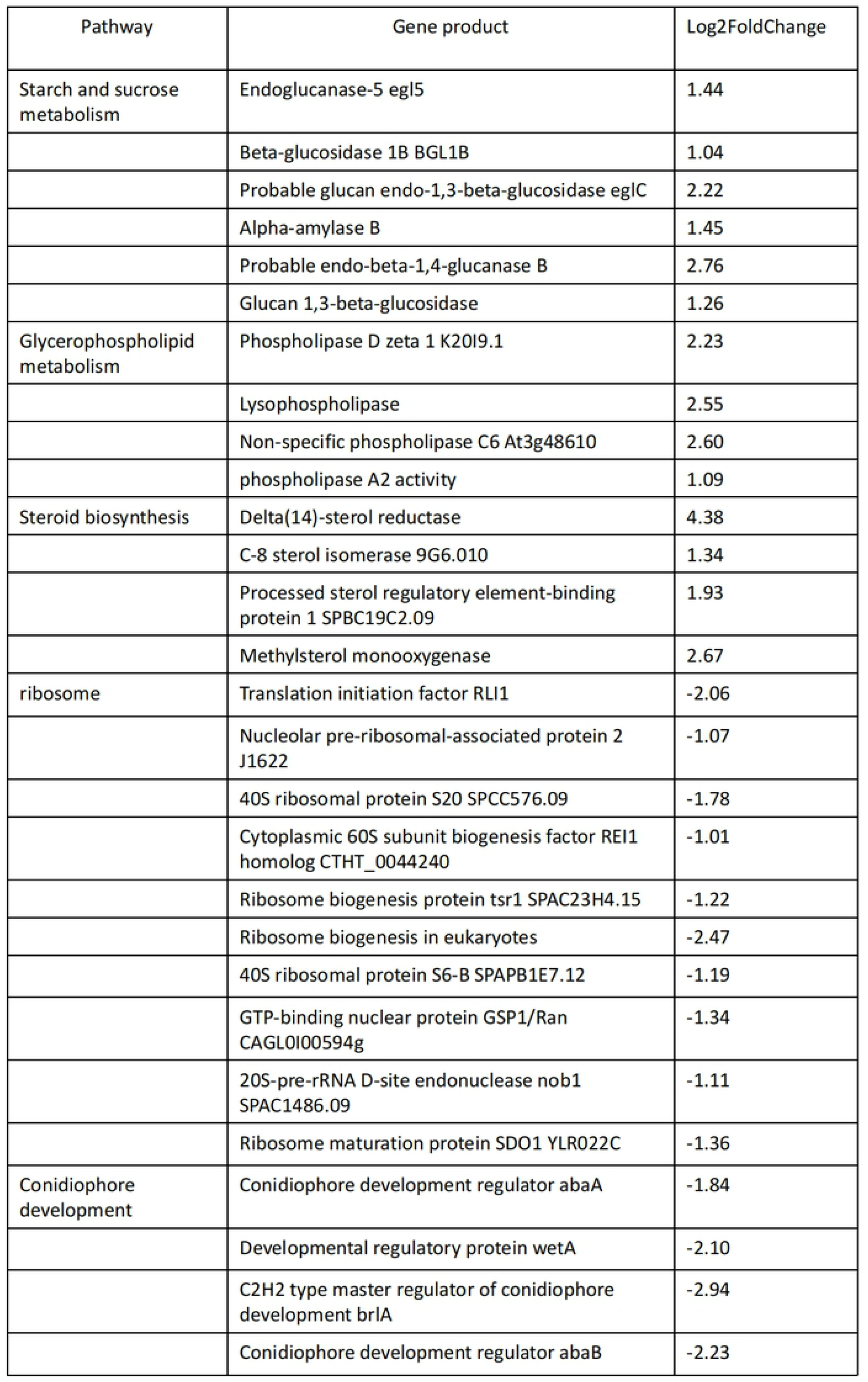
DEGs that may be involved in **HK** antifungal mechanism.

### In vitro and in vivo toxicity evaluation of HK

In vitro, HCECs were incubated with different concentrations of HK solution. CCK-8 experiment showed that HK was not toxic to HCECs in the concentration range of 2-12 μg/ml (Fig. 7A), of which 2 μg/ml and 4 μg/ml had the ability to promote proliferation. While the concentration of more than 12μg/ml HK solution reduced the proliferation ability of HCECs. In addition, wound healing showed that HK at 4-12 μg/ml did not affect migration ability of HCECs within 12 hours compared with the control group, while over 12 μg/ml HK reduced migration ability (Fig. 7B, C). In vivo, toxicity of HK was assessed using the Draize test by sodium fluorescein staining. No fluorescein sodium staining was seen in mouse corneas treated with DMSO or 10, 16 and 32 μg/ml HK for 1 day, 3 days and 5 days (Fig. 7B).

**Fig 7.**
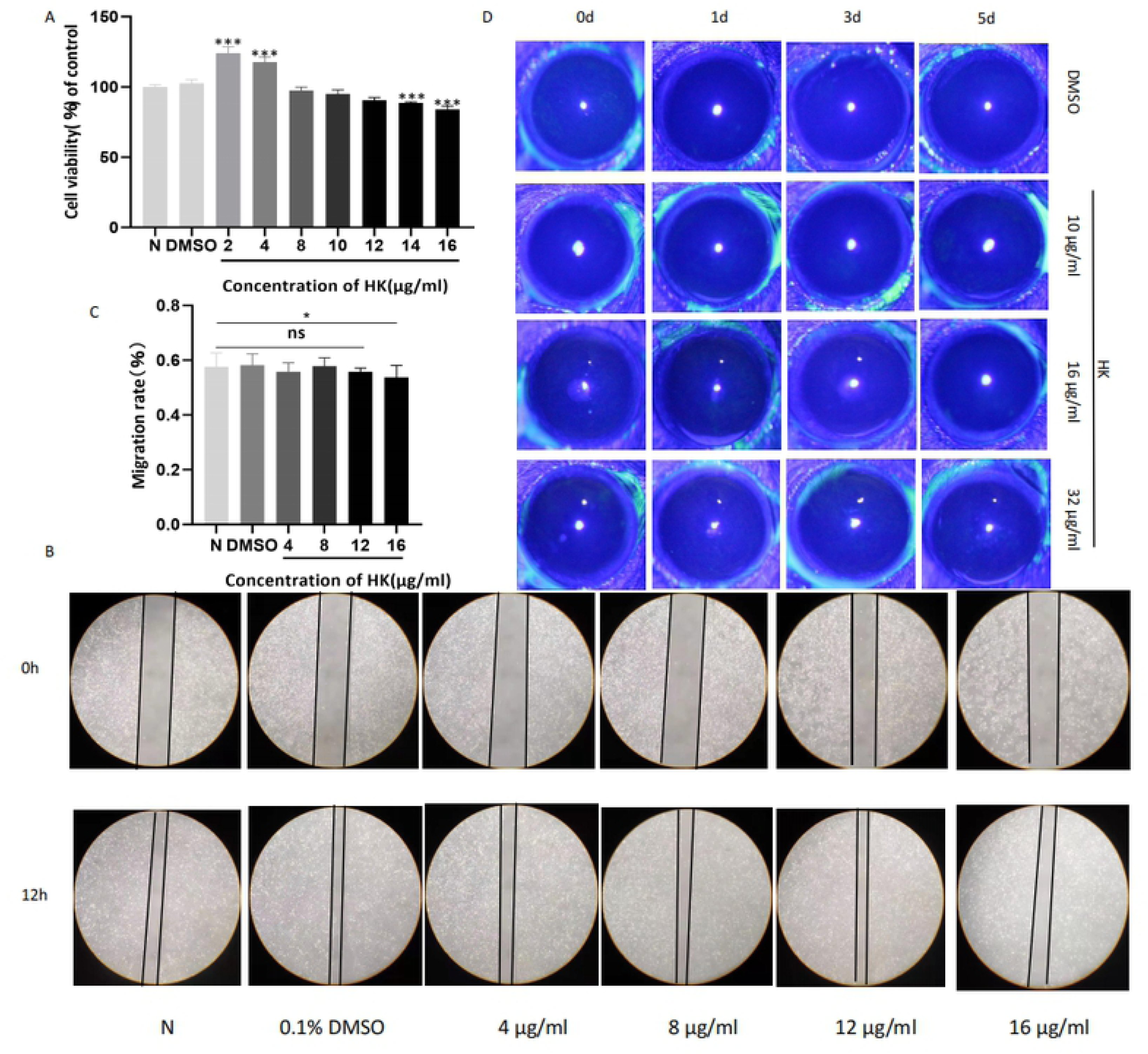
Effects of **HK** on cell viability and cornea toxicity. HCECs were treated with HK (0, 2, 4, 8, JO, 12, 14 and 16 µg/mL) or 0. J¾DMSO for 24 hours and then incubated with CCK-8 for 3 hours to explore the effect of HK on cell viability (A). Wound healing assey (B) and quantitative analysis (C) were used to evaluate the effect of HK on cell migration. The Draize Test was used to test the potential adverse effects of HK on cornea (D).

### HK eased murine FK

The effect of HK on FK was examined by slit-lamp photography and clinical score(24). Slit-lamp images showed that the HK-treated group had less corneal opacity, reduced ulcer area, and a lower clinical score on day 3 post-infection compared with the DMSO-treated cornea (Fig. 8A, B). The fungal burden was measured by fungal plate count. Compared to the DMSO-treated group, HK treatment reduced the number of viable fungal colonies in infected corneas (Fig. 8C, D).

**Fig 8.**
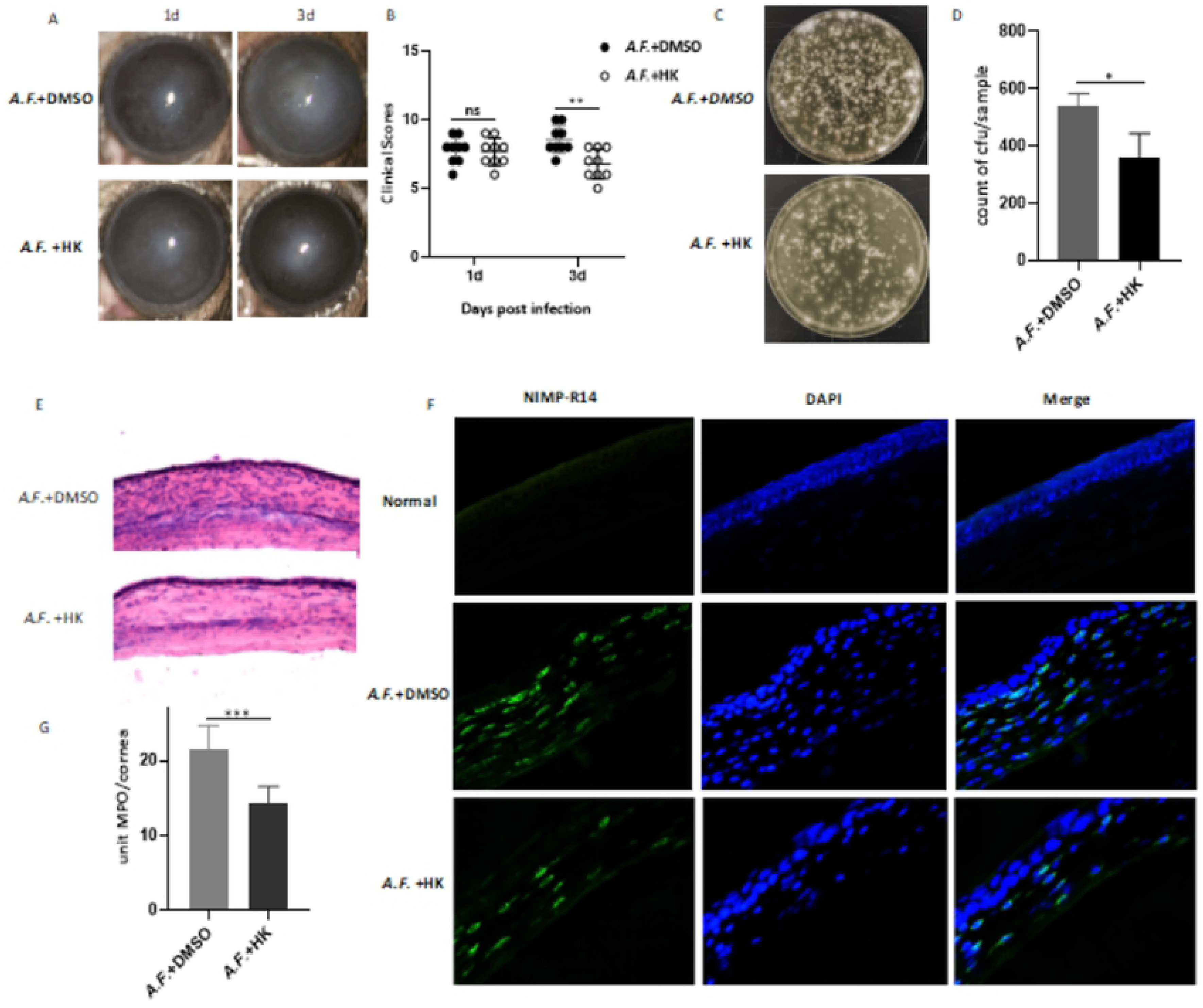
FK severity was alleviated followed by **HK** treatment compared with the DMSO group. The slit-lamp photography (A) and clinical score (B) showed that HK reduced corneal opacity and clinical scores on 3 days post-infection compared with the DMSO-treated cornea. HK treatment of C57BU6 mice reduced the number of viable fungal colonies in infected corneas compared with DMSO group (C) and the quantitative analysis (0). On 3 days post-infection, representative images of HE staining of the cornea treated with HK or DMSO (E, 400x). The fluorescence of neutrophils in corneas of normal group, HK-treatment group or DMSO-treatment group (F, 400x). The neutrophils were stained with NIMP-Rl4 with green fluorescence, and the nucleus is stained with DAPI and has blue fluorescence. MPO levels were shown in figure G.

HE staining of corneal tissue showed that on the 3rd day after infection, a large number of inflammatory cells were infiltrated in the corneal epithelial layer and stromal layer of the control group. While in HK group, the number of inflammatory cells was significantly reduced (Fig 8E). The localization and quantification of neutrophils in infected corneas were assessed by IFS and MPO assay. There were no neutrophils in the corneal stroma of normal mice, while the fluorescence of neutrophils increased in infected corneas. The fluorescence of neutrophils in HK group was significantly less than that in DMSO group (Fig. 8F). Similarly, the MPO level of the HK group was significantly decreased on the 3rd day after infection compared with the DMSO group (Fig. 8G).

To explore the anti-inflammatory mechanism of HK in murine FK, we detected the mRNA and protein expressions of LOX-1 and pro-inflammatory factors in infected corneal tissue. Compared with the control group, HK significantly inhibited *A.F.*-induced LOX-1 expression both at the mRNA level (Fig. 9A) and protein level (Fig. 9B, C). In addition, the mRNA levels of pro-inflammatory factors IL-1β, IL-6 and TNF-α in HK group were significantly lower than those in control group on 3rd day (Fig 9D, 9E, 9F). ELISA results showed that HK could significantly inhibit the protein expression levels of IL-1β, IL-6 and TNF-α (Fig 9G, 9H, 9I).

**Fig 9.**
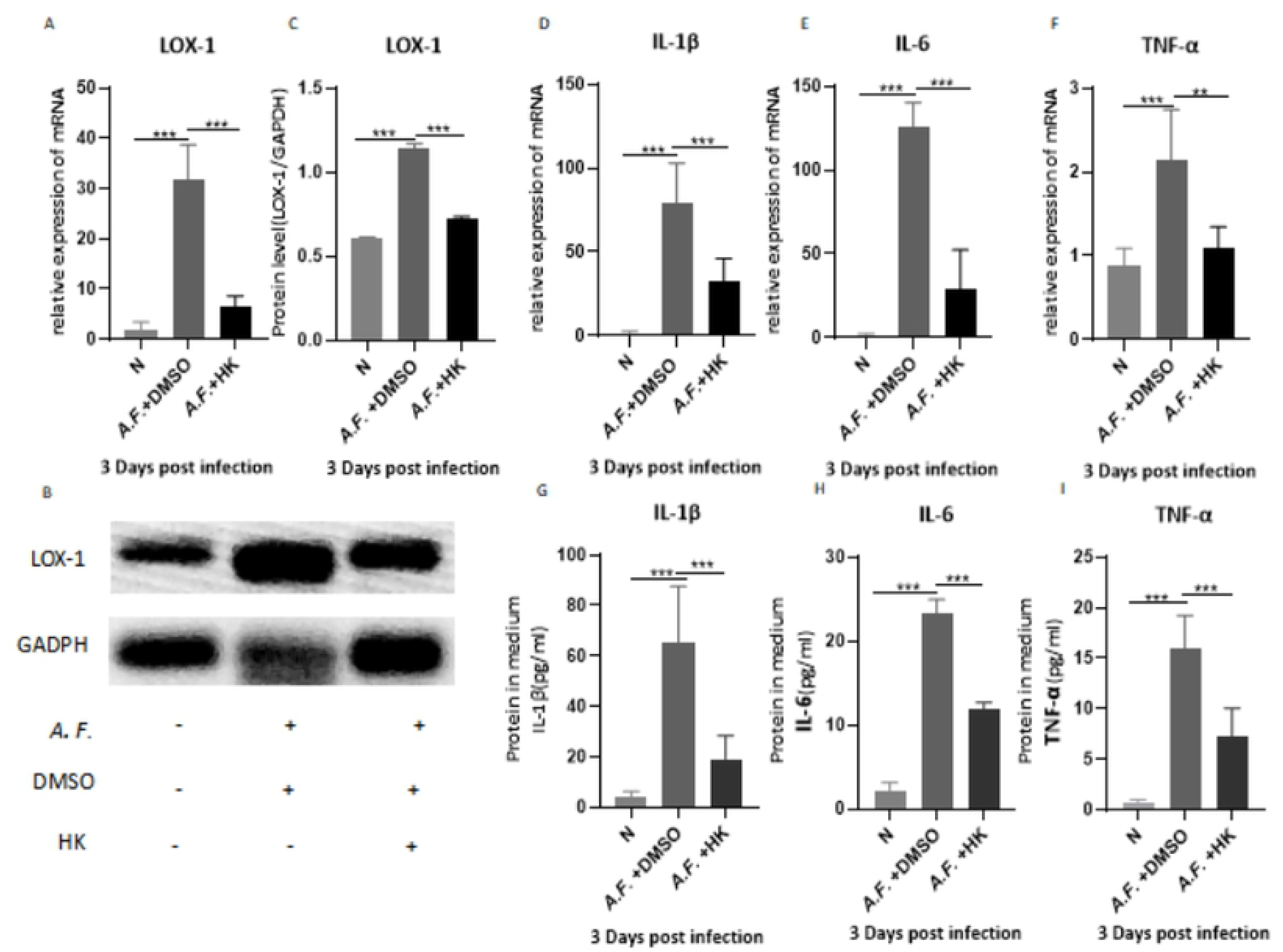
H**K** decreased the inOammatory mediators in corneas induced by *A.F..* qRT-PCR results for LOX-I (A), IL-IP (D), IL-6 (E), and TNF-u (F) in *A.J;:* infected mouse cornea treated with DMSO or HK 3 days post infection. The protein expression of LOX-I was detected by Western Blot (B, C). The protein levels of IL-Ip (G), IL-6 (H), and TNF-u (I) were tested by ELISA.

## Discussion

FK is a highly blinding infectious eye disease caused by fungal infections(25). After the fungus invades the corneal tissue, it causes an uncontrollable excessive inflammatory response, resulting in severe corneal injury, hyphema, and even corneal perforation(3, 26, 27).HK is a natural component found in *Chamaecyparis taiwanensis*, which has been found to have outstanding antifungal and anti-inflammatory effects(28). In the present study, we found that HK could not only effectively inhibit the growth of *A.F.*, but also reduce the inflammatory response of keratitis.

In vitro, HK inhibited the growth of *A.F.* in a concentration-dependent manner and significantly inhibited the growth of A.F. at 8 μg/ ml. Our result is similar to the recent study showing the growth inhibitory effect of HK on *C. albicans* at 5 μg/ml(11). SEM results showed that “pits” and “holes” which can result in cell wall and cell membrane rupture, were formed on the surface of HK-treated *A.F.* cells. In PI staining experiment, we found that the permeability of A.F. membrane was increased after HK treatment. Studies have shown that microorganisms can die due to damage to cell walls and membranes(29). The treated cells exhibited complete separation of the cell membrane from their wall, disorganized internal structure of the cell, and loss of some organelles. It is the first time to ultrastructurally identify the antifungal mechanism of HK. In addition to mycelium growth, adhesive force to corneal cells and biofilm formation of fungi are also important steps in infection(30). Biofilms are composed of polysaccharides, melanin, proteins and extracellular DNA and biofilm formation is critical step in fungal infection of host cells and resistance to the host immune system (31, 32). Our study confirmed that HK could inhibit *A.F*. adhesion and biofilm formation. Consistently, Kim et al. found that HK effectively inhibited the production of *C. albicans* biofilms. (12) These results suggest that HK can inhibit the adhesion ability and the biofilm formation of *A.F.*, thus reducing corneal fungal infection.

In order to explore the transcriptome changes of *A.F.* after treated with HK, GO and KEGG enrichment analysis was used after sequencing the transcriptomes. As essential components of the cytoskeleton, the expression of metabolism-related genes of carbohydrates and various amino acids was reduced, while the expression of lipid metabolism-related genes was increased, suggested that the cell cycle was disturbed by HK(33). It is known that starch metabolism and sucrose metabolism are involved in cell wall integrity.(34) Glucan 1,3-β-glucosidase catalyzes the hydrolysis of glucan, one of the main components of fungal cell walls.(35) Endo-β-1,4-glucanases and β-glucosidases are involved in the bio-degradation of cellulose, a major structural polymer in fungal wall(36). Sequencing results showed that HK treatment up-regulated the expression of glucan and cellulose hydrolysis-related enzymes (Table 2), suggested that HK may disrupt cell wall structure by inducing the gene expression of hydrolase and further increasing the sensitivity of fungal cells to HK(37).The expression levels of DEGs, related to membrane homeostasis-related pathways, such as glycerophospholipid metabolism and steroid biosynthesis, were significantly up-regulated after HK treatment (Table 2)(38). Phospholipases A and D are enzymes that catalyze the decomposition of phospholipids, the main components of cell membranes. After HK treatment, genes of phospholipases A and D were activated, suggested that HK may disrupt membrane integrity by altering membrane -related genes.

Furthermore, HK treatment down-regulated most of the DEGs associated with ribosome biogenesis of *A.F.* (Table 2). As the site of making protein, ribosome biogenesis is a complex process closely related to growth rate.(39) It has been shown that impaired ribosome disrupts the integration between morphogenesis and nucleus replication during fungal germination,(40) which is consistent with the morphological changes of hyphae after HK exposure observed by electron microscopy in our research. It was observed that genes related to genetic information processing-translation mainly decreased after HK treatment, suggesting that HK may exert its antifungal effect by inhibiting ribosome biosynthesis and subsequent protein biosynthesis. Interestingly, we found that from the genomic data genes related to *A.F.* asexual reproduction were down-regulated in HK group (Table 2). This was consistent with our qRT-PCR results that abaA, wetA and brlA were down-regulated after HK treatment. AbaA, wetA and brlA, their sequential activation is required during fungal asexual reproduction(41). Deletion of wetA leads to delayed germ-tube formation from conidia, and reduced density of hyphal branching and thallic(42). This change was also observed after treatment with different concentration HK, indicating the antifungal effect caused by the inhibiting of growth and reproduction of *A.F.*.

HK significantly relieved the corneal damage in our study, in which the antifungal effect of HK was first actually proven in murine fungal keratitis. Neutrophil is important immune cells in the pathogenesis of fungal keratitis(43), however excessive neutrophil infiltration with a large collection of active oxygen, free radical and lysosomal enzymes, often leads to severe damage to corneal tissue (44–46). HK treatment significantly reduced MPO levels and the depth of neutrophil infiltration in murine fungal keratitis corneas. Previous studies have found that in *Streptococcus pneumoniae* pneumonia, HK can lighten the inflammatory damage by reducing the infiltration of neutrophils and the expression level of inflammatory factors in the lung(47). The above results suggest that HK can play an anti-inflammatory role by inhibiting the neutrophil trafficking during inflammation.

Lee et al. found that HK could down-regulate the inflammatory response mediated by LPS through inhibiting the expression level of TNF-α and IL-6 in primary human keratinocytes(48). Our data indicated that HK also significantly down-regulated the mRNA and protein levels of corneal pro-inflammatory cytokines TNF-α, IL-6, and IL-1β, which could mediate the recruitment, activation and adhesion of neutrophils(49). A plentiful supply of virulence factors produced by the inflammatory cascade can damage the corneal epithelium and corneal stromal structure(50, 51). Lu et al. found that HK effectively prevented liver injury after hemorrhagic shock and resuscitation (HS/R) by inhibiting the inflammatory response(52), which is consistent with our findings. Our experiment also found that HK inhibited the expression of LOX-1 induced by *A.F..* Previous studies have shown that LOX-1 is an important pattern recognition receptor involved in corneal antifungal immune responses(22). Inhibition of LOX-1 results in the lower intracellular ROS production, p38-MAPK dephosphorylation, NF-κB translocation and aberrant BCl-2 expression, and down-regulate inflammation together(53). All results suggested that the down-regulated inflammatory response was associated with the decreased LOX-1 signaling.

In summary, we found that HK can exert antifungal effects by inhibiting the growth, adhesion, and biofilm formation of *A.F.*, as well as inducing fungal morphological changes, increasing the permeability of fungal membranes, and destroying the intracellular structure. HK interfered with the cell cycle at the transcriptional level, affecting cytoskeleton and ribosome synthesis. In addition, HK exerted a protective effect in *A.F.* keratitis, and its mechanism is related to that HK reduced corneal fungal burden, inhibited the expression of LOX-1 and pro-inflammatory factors, and reduced neutrophil infiltration.

Our work uncoverd a novel, remarkably powerful role for HK against *A.F.* and murine inflammatory response, suggesting HK has the potential to become a new treatment for FK. Future work will validate the signling targets of HK in more fungal pathogens and explore whether the fungistatic activity it possesses can be clinically adapted. In time, this may hold promise for the development of HK as antifungal agents in fungus-causing ocular inflammation.

## Materials and Methods

### HK Solution Preparation

HK powder (CAS 499-44-5), purchased from Sigma-Aldrich (Shanghai, China), was dissolved in DMSO to a storage concentration of 32 mg/ml, and diluted to suitable working solutions to achieve various final concentrations.

### *A.F.* culture

The standard *A.F.* strain (CPCC 3.0772), purchased from China General Microbiological Culture Collection Center, was cultivated on sabouraud agar medium for 2-3 days, and then conidia on the surface of the medium were gathered. Adjust the final concentration of 1×10^7^ CFU/ml in PBS with a hemocytometer.

### MIC for *A.F.* Conidia

MIC for *A.F.* conidia of HK was assayed by a standardized microdilution method in the 96-well plate described as before(54). *A.F.* conidia were prepared as described above. 32 μg/ml of HK in sabouraud medium was diluted to 6 different concentrations by two-fold gradient dilution, then transferred into third to eighth column wells (100 μL per well). The first column was the blank control, the second column was incubated with sabouraud medium with 0.1% DMSO. Finally, 5 μL prepared conidia suspension (1×10^7^ CFU/ml) was added into the 96-well plate. The plates were incubated at 28°C without shaking for 36 hours. *A.F.* conidia MIC was determined spectrophotometrically at 620 nm. Then, the supernatant in 96-well plates were discarded, and 50 μl of Calcofluor White Stain (Sigma-Aldrich, Shanghai, China) was added for 10 min at room temperature. The staining pictures were photographed using the fluorescence microscope (Nikon, Tokyo, Japan, 100×).

### MIC for *A.F.* Hyphae

Conidia suspension (5×10^5^ CFU/ml) was incubated in 96-well plates (100 μL per well) at 28 °C for 24 hours to form hyphae, then 0.1% DMSO was added with 2, 4, 8, 12, 16 and 32 μg/ml HK per well and incubated for 24 hours. *A.F.* hyphae MIC was determined spectrophotometrically at 620 nm.

### Time Kill Assay

Based on the MIC for *A.F.* conidia, the preparation of the microbial inoculum was carried out by the standardized microdilution method as described above. The absorbance at 24 hours, 48 hours, 72 hours and 96 hours was measured. The absorbance at 620 nm of each well represented the amount of precipitation in every well.

### SEM

*A.F.* conidia (1 × 10^7^ CFU/ml) was cultured in 6-well plates (1 ml per well) at 28°C for 24 hours to form hyphae. Then, the hyphae were washed, centrifuged (12,000 rpm, 10 minutes), and transferred to a new 6-well plate, followed by incubation with 0.1% DMSO or 2 μg/ml HK for 24 hours at 28°C. After PBS rinsing was performed three times, the hyphae were collected, fixed by 2.5% glutaraldehyde at 4°C for two hours. The samples were washed with PBS, then mixed with 1% (v/v) osmium tetroxide in PBS at 4°C for one hour. Subsequently, samples were gently dehydrated in graded ethanol, criticalpoint dried in CO_2_, coated with gold, and observed under SEM (VEGA3; TESCAN Company, Shanghai, China) at magnification × 2000 and × 5000 (bar = 20 or 10 μm).

### Propidium Iodide (PI) Uptake Testing

The conidia suspension of *A.F.* (1×10^7^ CFU/ml) was seeded into 12-well plates and incubated at 28°C for 24 hours to form hyphae. Then the hyphae were washed, centrifuged (12,000 rpm, 10 minutes), and transferred to a new 12-well plate, followed by incubation with 0.1% DMSO or HK (2, 4 μg/ml) for 24 hours at 28°C. After rinsing with PBS, 1 ml 50 μg/ml PI solution (Leagene biotechnology, Beijing, China) was added to each well for 15 minutes’ incubation at room temperature in the dark. Images were captured with a fluorescence microscope (Nikon, Tokyo, Japan, 100×) under green excitation light.

### TEM

The *A.F.* conidia (1×10^7^ CFU/ml) was seeded into 6-well plates and incubated at 28℃ for 24 hours to form hyphae. Then the hyphae were washed, centrifuged (12,000 rpm, 10 minutes), and transferred to a new 6-well plate, followed by incubation with 0.1% DMSO or HK (4 μg/ml) for 24 hours at 28℃. After rinsing with PBS, the hyphae were collected, fixed by 2.5% glutaraldehyde at 4℃ overnight. Experimental procedures for preparation of hyphae sample observed by TEM were performed according to routine methods(55). The samples were then observed in the JEOL-1200EX transmission electron microscope (JEOL Ltd., Tokyo, Japan).

### Biofilm Assay

Crystal violet assay was used to determine the biofilm forming capacity(56). Briefly, the preparation of the microbial inoculum before the biofilm formation was carried out by the standardized microdilution method as described on the MIC for *A.F.* conidia. After 48 hours incubation, biofilms were rinsed for 3 times, dried in air and fixed with 99% methanol for 20 minutes, and then stained with 0.1% crystal violet (Sigma-Aldrich, Shanghai, China) for 15 minutes. PBS washed out unbound dyes until the eluent is colorless. After air drying, 100 μ L 95% ethanol was added to each well at room temperature for 30 minutes to fully release the dye combined with the biofilms. The supernatant was transferred to a new 96-well plate and the OD value of each well was measured at 570 nm three times.

### Fungal Adherence Assay

Conidia suspension (2 × 10^5^ CFU/ml) containing 0.1%DMSO or HK (8 μg/ml) was mixed with HCECs (provided by Laboratory, University of Xiamen, Fujian, China) (2 × 10^4^/ml) and plated on the chambered slides (4/slide) as described previously(26). Each slide was incubated at 37°C for 3 hours, then washed with sterile PBS and stained by hematoxylin and eosin (HE) staining. The spores adhering to HCECs were observed and photographed by an optical microscopy (Nikon, Tokyo, Japan, 400×).

### Sample Preparation for RNA-Seq

Conidia suspension (2 × 10^5^ CFU/ml) containing 0.1%DMSO or HK (2 μ g / ml) was inoculated on a 6-well plate at 28 °C. The Con (DMSO treated) and Test (HK treated) samples were harvested after incubation for 36 hours, and then stored at 80 °C for RNA-Seq analysis. All experiments were carried out in 3 independent biological replicates.

### RNA Isolation and Library Preparation

Total RNA was extracted using the TRIzol reagent according to the manufacturer’s protocol. RNA purity and quantification were evaluated using the NanoDrop 2000 spectrophotometer (Thermo Scientific, USA). RNA integrity was assessed using the Agilent 2100 Bioanalyzer (Agilent Technologies, Santa Clara, CA, USA). Then the libraries were constructed using TruSeq Stranded mRNA LT Sample Prep Kit (Illumina, San Diego, CA, USA) according to the manufacturer’s instructions. The transcriptome sequencing and analysis were conducted by OE Biotech Co., Ltd. (Shanghai, China).

### RNA-Seq and DEGs Analysis

The libraries were sequenced on an Illumina HiSeq X Ten platform and 150 bp paired-end reads were generated. Raw reads for each sample were generated. Raw data (raw reads) of fastq format were firstly processed using Trimmomatic and the low-quality reads were removed to obtain the clean reads(57). Then clean reads for each sample were retained for subsequent analyses.

The clean reads were mapped to the *A.F.* genome (NCBI_ASM265v1) using HISAT2(58). FPKM of each gene was calculated using Cufflinks, and the read counts of each gene were obtained by HTSeq-count(59). Differential expression analysis was performed using the DESeq (2012) R package. P value < 0.05 and foldchange > 2 was set as the threshold for significantly differential expression. Hierarchical cluster analysis of differentially expressed genes (DEGs) was performed to demonstrate the expression pattern of genes in different groups and samples. GO enrichment and KEGG pathway enrichment analysis of DEGs were performed respectively using R based on the hypergeometric distribution(60).

### Quantitative real-time Polymerase Chain Reaction (qRT-PCR) verification

To validate the RNA-seq gene expression patterns, nine differentially expressed genes, with a putative function relative to the conidium formation, cell wall formation, were selected to confirm the RNA-Seq data by qRT-PCR. The total RNA was prepared using the Fungal Total RNA Isolation Kit (B518629, Sangon Biotech, Shanghai, China), and reverse transcribed with HiScript III RT SuperMix (Vazyme, Nanjing, China) according to the manufacturer’s instructions. The PCR method was based on previous studies(61). Primers used for the qRT-PCR are listed in S1 Table.

### Cell Viability Assay (CCK-8)

HCECs were suspended and inoculated in the 96-well plate and treated with HK (0, 2, 4, 8, 10, 12, 14 and 16 μg/ml) or 0.1%DMSO for 24 hours. The cells were incubated for 3 hours with Cell Counting Kit-8 (CCK-8; MCE, New Jersey, America), and the absorbance was measured at 450 nm. Each sample had six replicates.

### The Draize Test

The potential adverse effects of HK in normal mouse eyes were tested by ocular toxicology study (Draize Eye Test). HK (10, 16, and 32 μg/ml) in dose of 5 μl was dropped into the conjunctival sac of one eye in four mice. The contralateral eye treated with 0.1%DMSO served as a control. Slit lamp microscopy under cobalt blue light was used to observe the corneal fluorescein staining (CFS) at 1, 3 and 5 days after corneal perfusion in mice(62).

### Wound healing

HCECs (3 × 10^5^ / ml) was seeded in a 6-well plate and incubated overnight at 37 °C. Then, three parallel lines were scratched on the cell layer using a 200 μl pipette tip in each well. HCECs was incubated with 0, 4, 8, 12 and 16 μg/ml HK or 0.1%DMSO for 12 hours. The width of cell scratches at the same location was measured at 0 and 12 hours under optical microscope (Nikon, Tokyo, Japan, 100×) to evaluate cell migration. The experiment was repeated at least three times under the same conditions.

### Murine Models of FK

Healthy C57BL/6 mice (female, 8 weeks old) were purchased from SPF (Beijing, China) Biotechnology Co., Ltd. All mouse treatments were in accordance with the ARVO Statement for the Use of Animals in Ophthalmic and Visual Research. The Ethics Committee on Experimental Animals approved our experiment and the approved number is: QYFYWZLL26777. Mice were abdominally anesthetized with 8% chloral hydrate. Then 2 μ L *A.F.* conidia suspension (1×10^7^ CFU/ml) was loaded into a sterile microsyringe (10 μL; Hamilton Corp., Bonaduz, GR, Switzerland) and inserted obliquely into the midstromal level in the center of the right cornea. The left eyes were blank control.

On the first day after modeling, the eyes in the HK group were treated with 5 μl HK eyedrop (10 μg/ml), and the eyes in control group were treated with 0.1%DMSO. These treatments were three times per day (once every 4 hours during the daytime). No treatment was given to the normal cornea. Slit lamp photography and clinical score (the standard of clinical scoring referred to Wu at al. was expressed in S2 Table(24)) were taken every day. The corneas of mice were removed by a scalpel and microscissor at the indicated time after treatment for qRT-PCR, Westernblot, myeloperoxidase (MPO), plate count and enzyme-linked immunosorbent assay (ELISA). The whole eyes were taken for immunohistochemical fluorescence staining (IFS) and HE Staining.

### Plate Count

The corneas of DMSO control group and HK treatment group were placed in PBS at 3 days p.i., then ground with a grinding stick and plated on Sabouraud agar mediums in triplicate. The plates were cultured overnight at 28 °C, and the number of visible fungal colonies on the plates was counted (n = 3/group/time) to reflect viable fungi surviving on the cornea(63).

### HE Staining

The eyeballs of mice were harvested at 3 days p.i. (n=3/group/time) and fixed with 4% paraformaldehyde at 4 °C for 3 days. After the lenses were removed, the eyeballs were embedded in paraffin and were filleted into 8 μ m under a cryostat. The sections were stained with hematoxylin and eosin and photographed under an optical microscope (Nikon, Tokyo, Japan, 400×).

### IFS

The eyeballs of mice were removed at 3 days p.i. (n=3/group/time), embedded in optimal cutting temperature (O.C.T) compound (Sakura Finetek USA, Inc., Torrance, CA, USA), frozen in liquid nitrogen. The 10 μm slices were fixed in acetone for 30minutes, then blocked with 10% goat serum (Solarbio, Shanghai, China) at room temperature for 30 minutes. Then these sections were incubated with rat anti-mouse NIMP-R14 antibody (Santa Cruz Biotechnology, Dallas, TX, USA) overnight at 4℃. After being washed with PBS, sections were stained with FITC-conjugated goat anti-rat secondary antibody (1:200; Elabscience) for 1 hour. The cell nuclei were stained with DAPI for 10 minutes. Images were captured by a fluorescence microscope (Nikon, Tokyo, Japan, 400×).

### MPO Assay

Mouse corneas (n = 6/group) were removed at 3 days p.i. and treated following the protocol of MPO kit (Nanjing Jiancheng Bioengineering Institute, Nanjing, China). The corneal tissue homogenate and reagent 3 were mixed at 9:1 and placed in water bath at 37 °C for 15 minutes. Add double distilled water, reagent 4 and chromogenic agent at 37 °C for 30 minutes. Add reagent 7, mix fully, and put it in water bath at 60 °C for 10 minutes. The absorbance was measured immediately at 460 nm.

### qRT-PCR on Cornea

Total RNA from mouse corneas was extracted by the RNAiso plus reagent (TaKaRa, Dalian, China). RNA samples were reverse transcribed using HiScript III RT SuperMix (Vazyme, Nanjing, China) to produce cDNA template. The PCR method was based on previous studies(61). The primer details are listed in S3 Table.

### Western Blot

Six corneas as a sample were lysed in 196 μl RIPA buffer, 2 μl PMSF and 2 μl phosphatase inhibitor for 2 hours. The protein concentration was determined by BCA assay (Solarbio, Beijing, China). The proteins were separated by SDS-PAGE electrophoresis and transferred onto polyvinylidene difluoride membrane (Solarbio). After blocking with blocking solution (Solarbio), the membrane was incubated with GADPH (1:2000; Elabscience, Wuhan, China) and LOX-1 (1: 1000: Abcam, Cambridge, MA, USA) at 4 °C. The membrane was washed in PBST for 3 times and then incubated with secondary antibody at 37 °C for 1 h. Then the spots are visualized with ECL Western blot detection reagents (Biotime, China).

### ELISA

Mouse corneas were collected and homogenated. The release levels of IL-1β, TNF-α and IL-6 in the supernatant were detected by mouse ELISA kit (Elabscience). The optical density of each hole was measured immediately at 450nm, and the reference wavelength was 570nm.

### Statistical analysis

*Mann-Whitney U* test was used to analyze the difference of clinical scores between the two groups at each time point. For comparisons with data from control group or HK treatment group, we performed an unpaired, two-tailed Student’s *t*-test or one-way *ANOVA* with post hoc analysis. GraphPad Prism 8.0 and ImageJ1.44p were used for statistical analyses, and values presented as the mean ± standard deviation (SD). *P* < 0.05 (\**P* < 0.05, \*\**P* < 0.01, \*\*\**P* < 0.001) was considered as statistically significant (ns = no significance). All experiments were repeated at least three times to ensure accuracy.

## Supporting information

**S1 Table.**
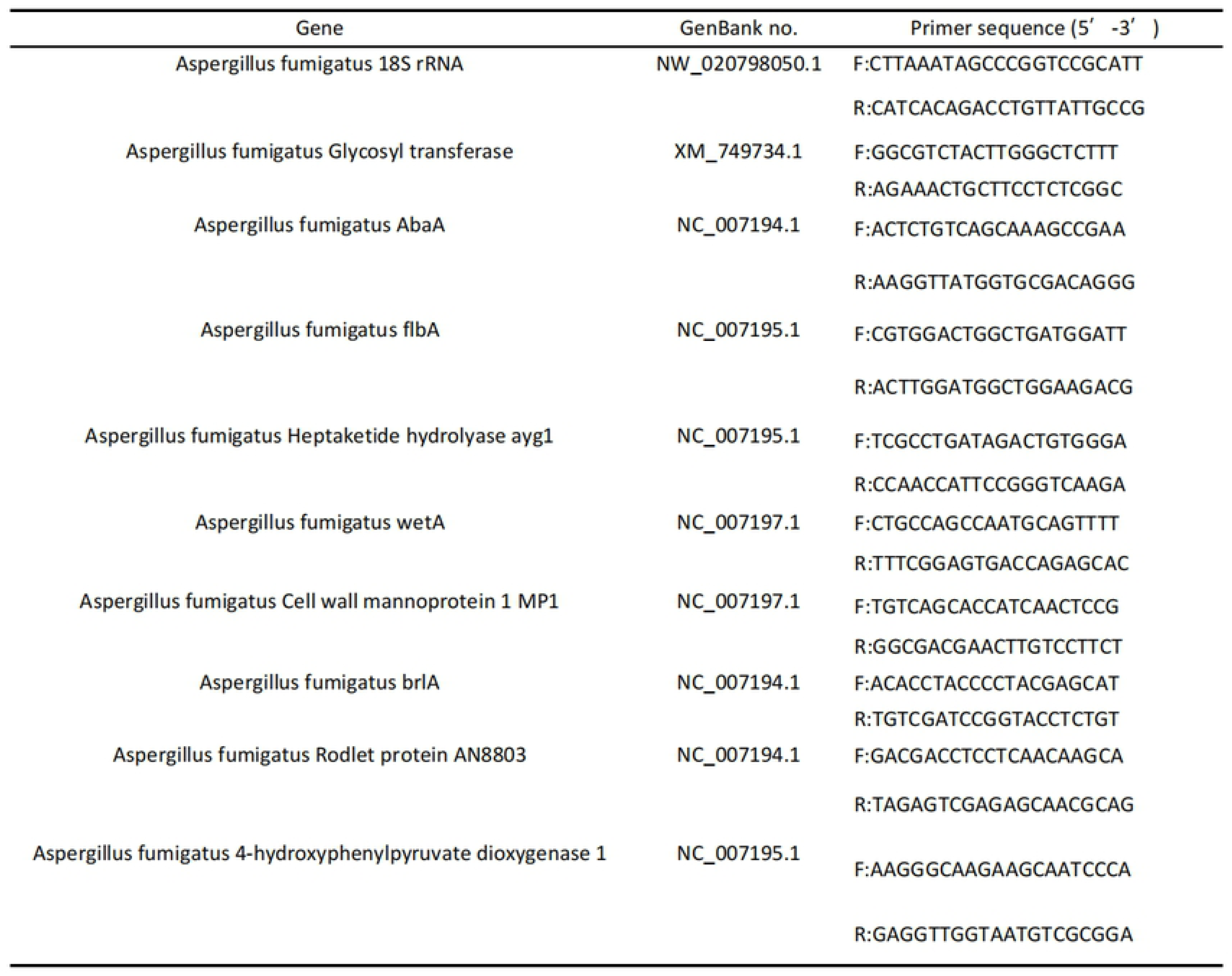
Primer List Used for *A.F.* qRT-PCR.

**S2 Table.**
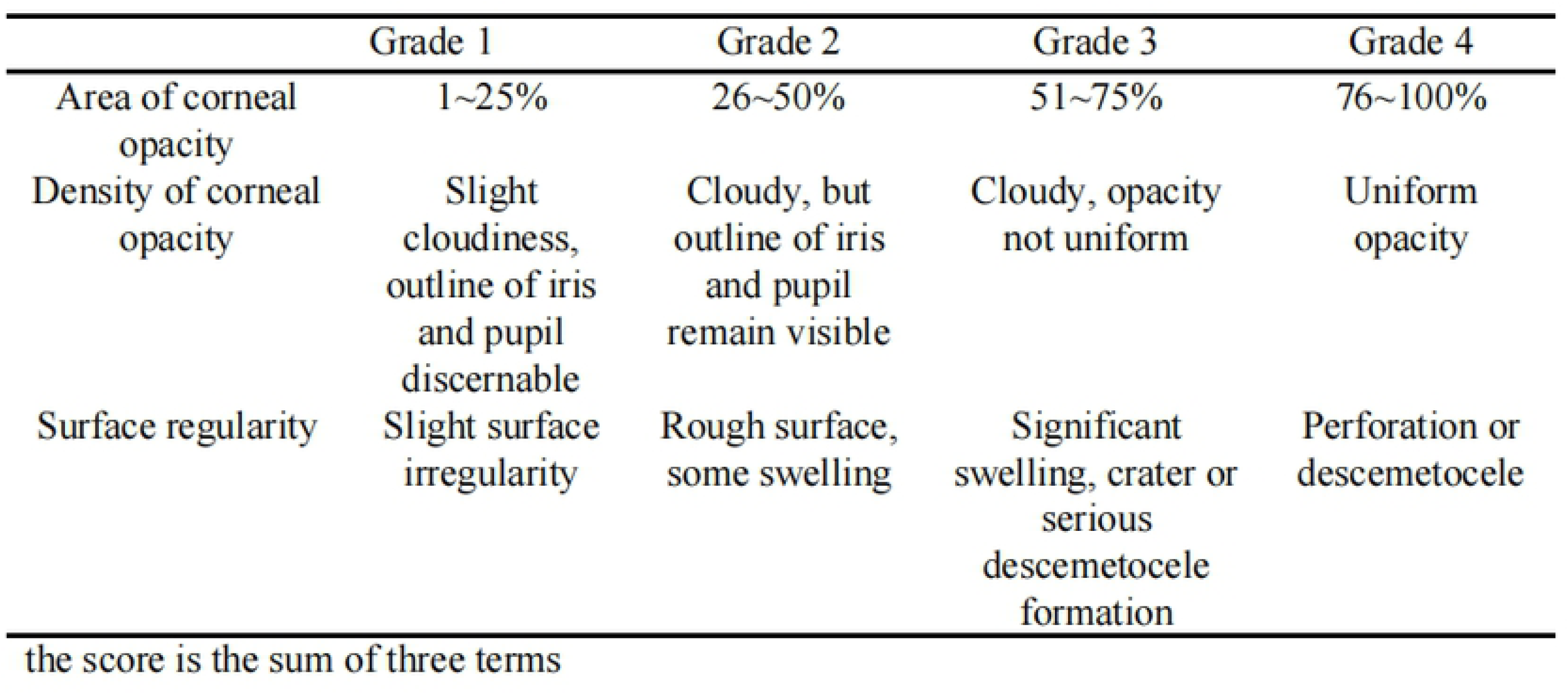
Visual scoring system for murine fungal keratitis.

Visual Scoring System for Murine Fungal Keratitis
|  | Grade 1 | Grade 2 | Grade 3 | Grade 4 |
| --- | --- | --- | --- | --- |
| Area of corneal opacity | 1~25% | 26~50% | 51~75% | 76~100% |
| Density of corneal opacity | Slight cloudiness, outline of iris and pupil discernable | Cloudy, but outline of iris and pupil remain visible | Cloudy, opacity not uniform | Uniform opacity |
| Surface regularity | Slight surface irregularity | Rough surface, some swelling | Significant swelling, crater or serious descemetocoele formation | Perforation or descemetocoele |

the score is the sum of three terms

**S3 Table.**
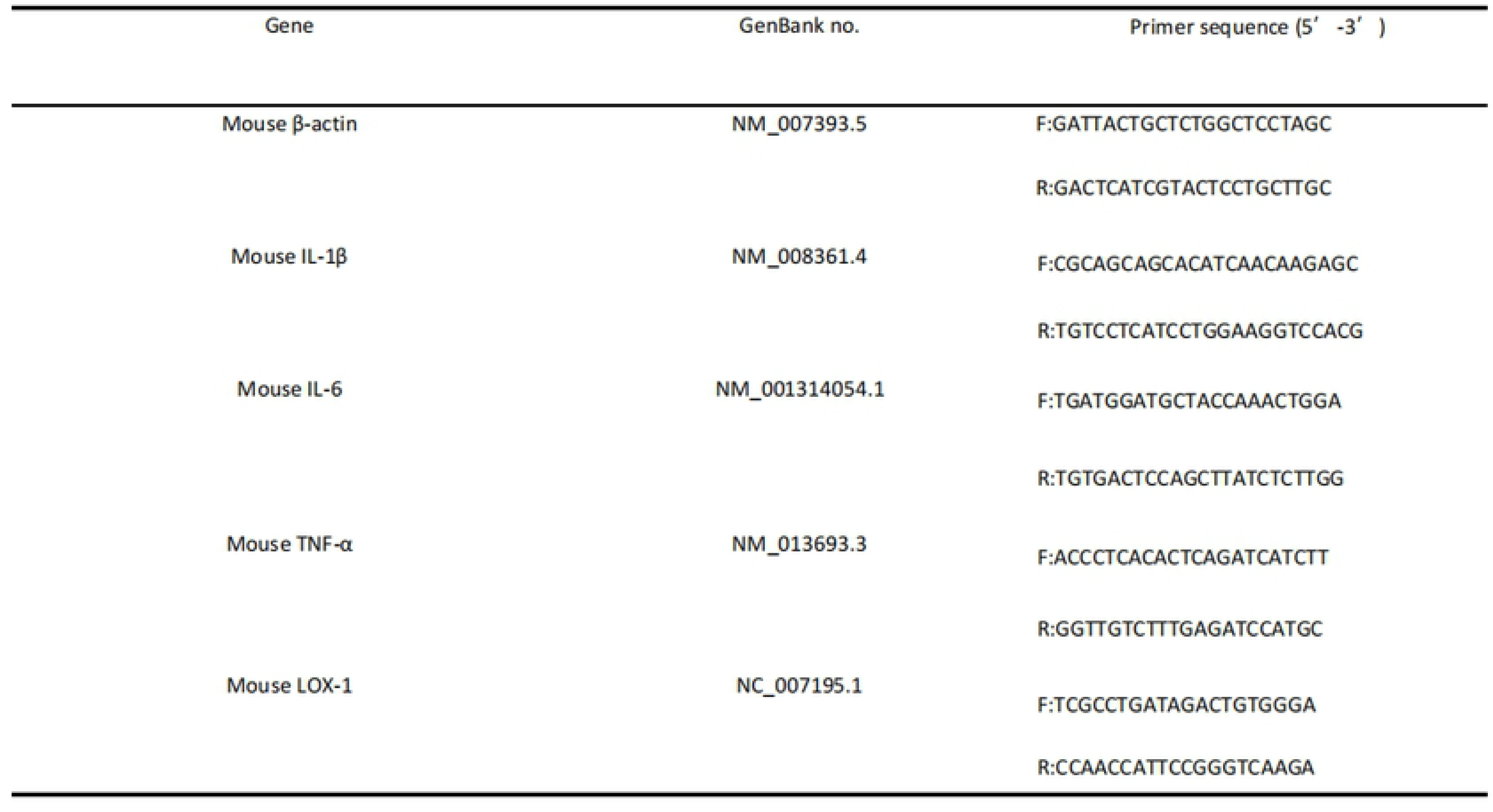
Primer List Used for Mouse cornea qRT-PCR.

## Acknowledgments

The content is solely the authors’ responsibility and does not necessarily represent the funding organizations’ official views.

